# Nano- and microplastics in pediatric tonsil tissue: bioaccumulation, distribution, and immunomodulatory effects in human lymphoid aggregate organoids

**DOI:** 10.64898/2026.05.27.728317

**Authors:** Amirbahador Golchin Sani Masouleh, Antony Georgiadis, Maggie Zhang, Yi-Wei Lin, Isha Kandlikar, Patrick Kiessling, Mitra Alkani, Andrea Miranda, Nathan Alves, Alex D. Bindemann, Anushri Umesh, Matthew Campen, Robert Taylor, Stacey Harper, Kara Meister

## Abstract

Nano- and microplastics (NMPs), by-products of the fragmentation and degradation of plastic products, are ubiquitous environmental contaminants, yet their burden in pediatric immune tissues and functional consequences for developing immunity remain unknown. Here we report the first comprehensive characterization of NMPs in surgically excised pediatric tonsils (n = 30) using pyrolysis gas chromatography– mass spectrometry (Py-GC/MS), Nile Red fluorescence microscopy, and optical photothermal infrared (O-PTIR) spectroscopy. NMPs were detected in all specimens, with polystyrene, polyethylene, polyethylene terephthalate, and acrylonitrile butadiene styrene present in >90% of samples. To bridge clinical exposure data with mechanistic insight, we formulated a cryo-milled multi-polymer mixture reflecting the patient-derived polymer profile and challenged human lymphoid aggregate culture (HLAC) tonsil organoids at environmentally relevant concentrations. Multiplexed cytokine profiling of culture supernatants revealed a robust early inflammatory response at day 3, with significant upregulation of IL-6 (p = 0.011) and MIP-1β/CCL4 (p = 0.011), followed by convergence toward control levels by day 14. Functional cytokine modules spanning immune, metabolic, structural, and growth factor pathways showed coordinated deviation from controls at day 3 post-exposure with subsequent normalization. Fluorescence-guided depth profiling demonstrated time-dependent penetration of 100 nm particles into organoid aggregates (70% tissue depth at day 3 versus 95% at day 14), and transmission electron microscopy revealed intracellular polyethylene within lymphocyte lysosomes. These findings establish pediatric tonsils as a sentinel tissue for NMP bioaccumulation and demonstrate that environmentally relevant polymer mixtures elicit transient but significant immunomodulatory responses in human lymphoid tissue, with implications for mucosal and systemic immune health in children.

Structure: Translational pipeline from clinical tissue characterization to patient-informed preclinical modeling of nano-microplastic (NMP) exposure in pediatric lymphoid tissue.
Pediatric tonsil tissue collected from clinically indicated tonsillectomies underwent tissue digestion for NMP characterization to identify NMP type and size distributions. In parallel, tonsil tissue was used to generate human lymphoid aggregate culture (HLAC) organoids that recapitulate the cellular complexity of the native tissue. These patient-derived organoids were then exposed to environmentally relevant compositions and concentrations of NMPs over time-course experiments, with longitudinal assessment of immunomodulatory responses including cytokine profiling and functional readouts. This bedside-to-bench approach establishes a physiologically relevant human system for investigating NMP-immune interactions, bridging clinical tissue analysis with mechanistic preclinical modeling to inform understanding of pediatric environmental exposures and their potential health impacts.

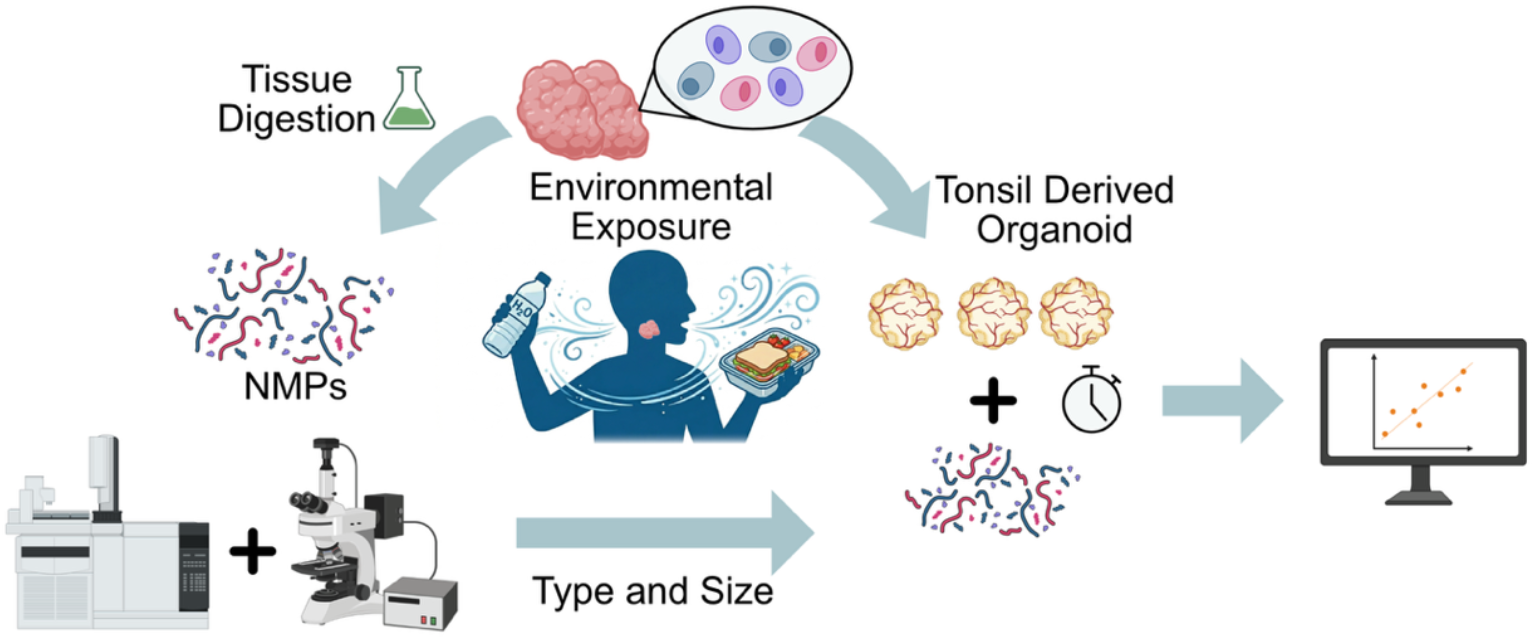

## Introduction

Global plastic production exceeds 430 million tons annually and continues to grow at approximately 4-5% per year.^1–3^ The resulting environmental burden of nano- and microplastics (NMPs) has become a public health concern.^4^ Nanoplastics are plastic particles smaller than 1 μm, while microplastics range in size from 1 μm to 5 mm.^5^ NMPs enter the human body primarily through ingestion and inhalation, and have been detected in blood, stool, lung, liver, brain, placenta, and other tissues.^6–9^

Recent clinical evidence has linked NMP accumulation to adverse cardiovascular outcomes, and experimental models demonstrate that NMPs cross epithelial barriers, reach systemic circulation, and accumulate in organs including the brain, where concentrations have increased significantly over time.^6^ Emerging evidence also links NMP accumulation to adverse digestive, reproductive, and respiratory health outcomes.^10,11^

Despite this growing body of evidence, critical knowledge gaps persist. First, there is a paucity of data on NMP burden in pediatric populations, who may be disproportionately vulnerable owing to higher respiratory rates, greater hand-to-mouth behavior, and developing immune systems.^12^ Also, since plastic production is increasing, pediatric populations may be exposed to higher concentrations at an earlier age than previous generations. Second, no study has systematically characterized NMP accumulation in human lymphoid tissue, responsible for orchestrating immune responses to environmental antigens. Third, mechanistic studies of NMP immunotoxicity have relied predominantly on isolated cell lines or animal models using single-polymer exposures at supraphysiological concentrations that were not informed by analogous human-tissue findings, limiting translational relevance.^12,13^

The palatine tonsils represent a uniquely informative tissue for addressing these gaps. As secondary lymphoid organs positioned at the gateway of the aerodigestive tract, tonsils are directly exposed to inhaled and ingested particles while simultaneously serving as immunological sentinels with dense lymphoid and vascular architecture, active germinal centers, and robust innate and adaptive immune capacity.^14^ In children, tonsils reach peak immunological activity between 3 and 10 years of age before gradual age-dependent involution.^15^ Tonsillectomy is one of the most common pediatric surgical procedures.^16^ It provides ethically accessible, paired tissue specimens that can be collected with minimal plastic instrument exposure. Furthermore, the tonsillar microenvironment harbors abundant bacteria implicated in plastic biodegradation, potentially facilitating *in situ* fragmentation of larger particles into nano-scale debris.^17^

Here we present an integrated clinical-preclinical investigation addressing two fundamental questions: (1) what NMPs are present in pediatric tonsil tissue, (2) how they might influence immune function. This research provides critical insights into compositional profiling and functional immunology of NMPs in pediatric tonsil tissue.

Most importantly, we use the clinical polymer profile found in tissue samples to directly inform preclinical exposure conditions. This bedside-to-bench approach creates, to our knowledge, the first patient-informed NMP substrate in a matched human organoid system.

## Results

### Ubiquitous NMP contamination of pediatric tonsils

Py-GC/MS analysis of pediatric tonsil specimens (n = 30) revealed NMP contamination in all samples tested. Three tissue preparation methods were compared: ethyl acetate/methanol extraction (n = 11), KOH with ultracentrifugation (n = 11), and standard KOH with cyclohexane filtration (n = 29). KOH with cyclohexane filtration method was selected because it was the method that achieved >65% probability for all polymers. After applying a >65% probability threshold for polymer identification, polyethylene (PE), polyethylene terephthalate (PET), and acrylonitrile butadiene styrene (ABS) were the most frequently detected polymers, present in >90% of samples (Fig. 1 a,b).

**Figure 1:**
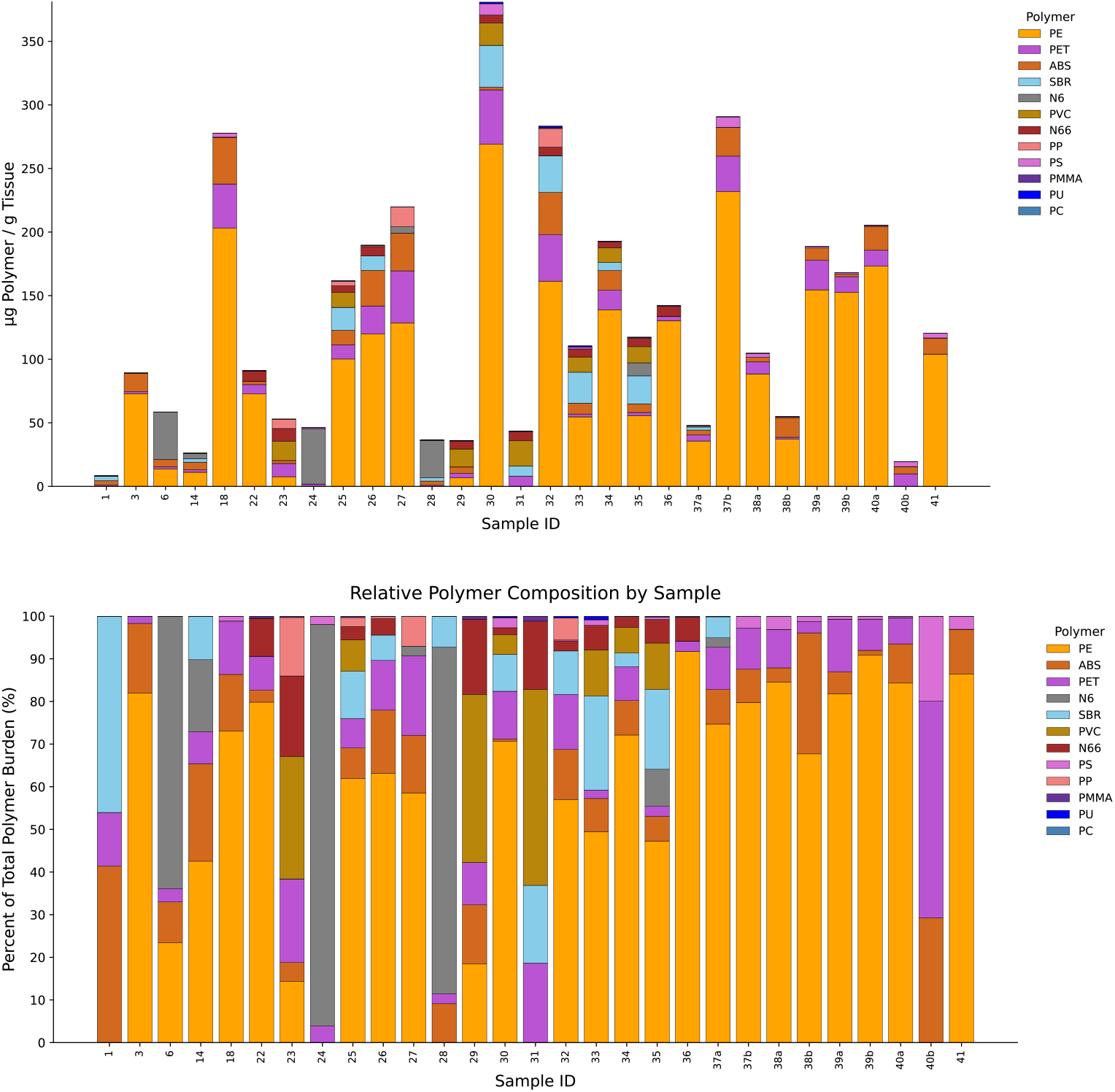
Ubiquitous NMP contamination of pediatric tonsils. A) Each sample showed muti-polymer signals. B) polyethylene (PE), polyethylene terephthalate (PET), and acrylonitrile butadiene styrene (ABS) were the most frequently detected polymers, present in >90% of samples. Additional polymers identified included polystyrene (PS), styrene-butadiene rubber (SBR), nylon-6 (N6), polyvinyl chloride (PVC), nylon-6,6 (N66), poly(methyl methacrylate) (PMMA), polycarbonate (PC), polyurethane (PU) and polypropylene (PP).

The polymer composition profile in pediatric tonsils showed notable differences from profiles reported in other human tissues. While PE predominates in brain and liver specimens, tonsils exhibited a more heterogeneous polymer distribution, consistent with their dual exposure to both inhaled and ingested particles.^6,7,9^ This finding aligns with recent systematic reviews demonstrating tissue-specific polymer accumulation patterns across human organs.

### Unsupervised clustering reveals distinct polymer exposure profiles

Principal component analysis (PCA) of polymer concentration profiles from KOH/filtration-processed tissue samples (n = 29) demonstrated separation of samples into two major clusters in multivariate space (Fig. 2). Prior to PCA, polymer concentrations were standardized by z-score normalization to account for differences in polymer abundance scales, and values below polymer-specific limits of quantification (LOQs) were excluded from analysis and set to zero in the feature matrix. The first two principal components explained approximately 54.5% of total variance (PC1: 31.7%; PC2: 22.8%). K-means clustering (k = 2) identified subgroups that separated primarily along PC1, suggesting differences in relative polymer composition rather than overall polymer burden alone.

**Figure 2.**
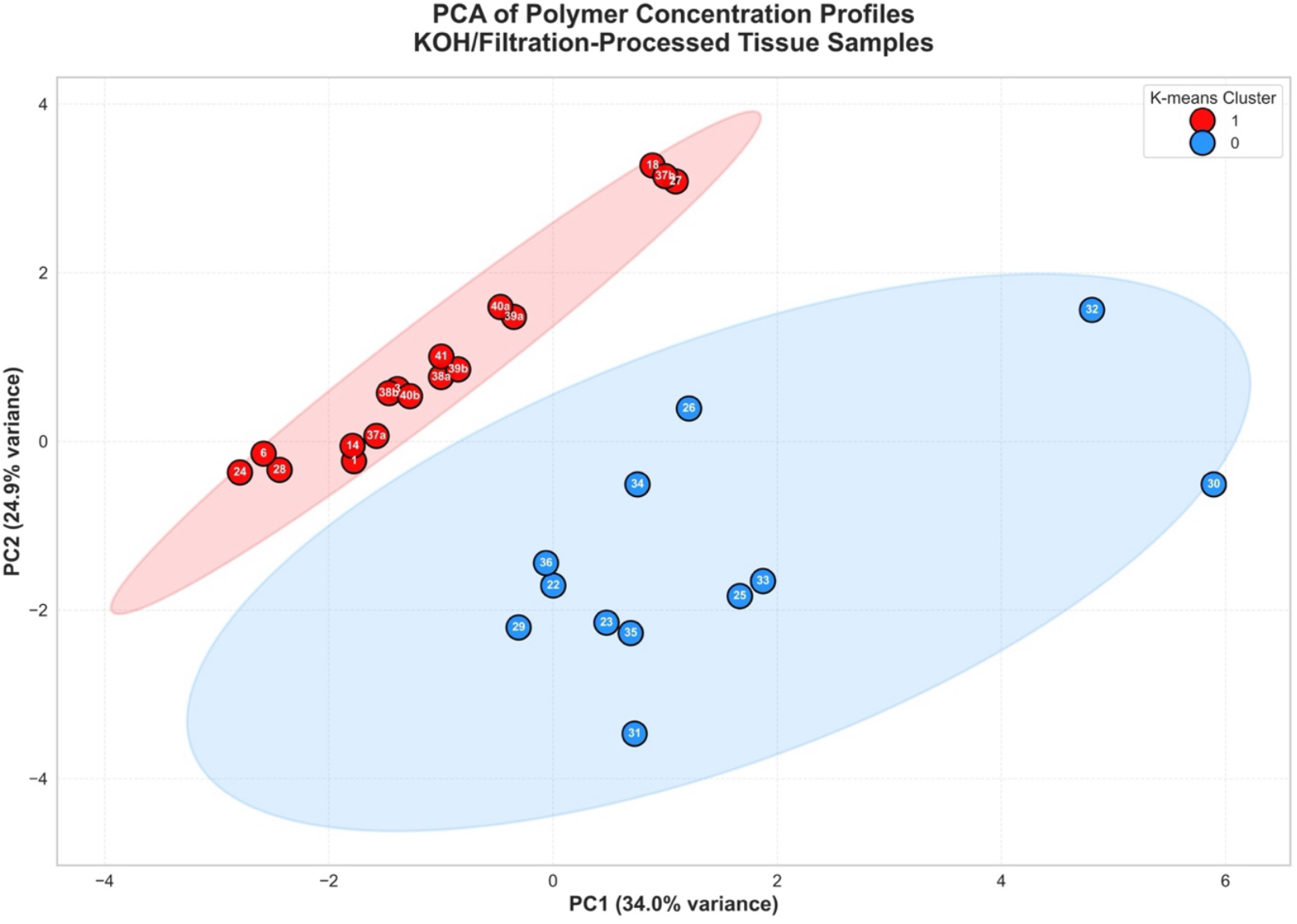
Principal component analysis (PCA) of polymer concentration profiles from KOH/filtration-processed tissue samples (n = 29). Polymer concentrations were z-score standardized prior to PCA, and missing or sub-LOQ polymer values were represented as 0 in the feature matrix. The first two principal components explained 58.9% of total variance (PC1: 34.0%; PC2: 24.9%). K-means clustering (k = 2) identified two major compositional groups that separated primarily along PC1. Points represent individual samples labeled by sample ID, and shaded ellipses represent approximate 95% confidence intervals for each cluster. Preliminary PCA loading analysis suggested that select polymers, including SBR, may contribute to cluster separation

**Figure 3:**
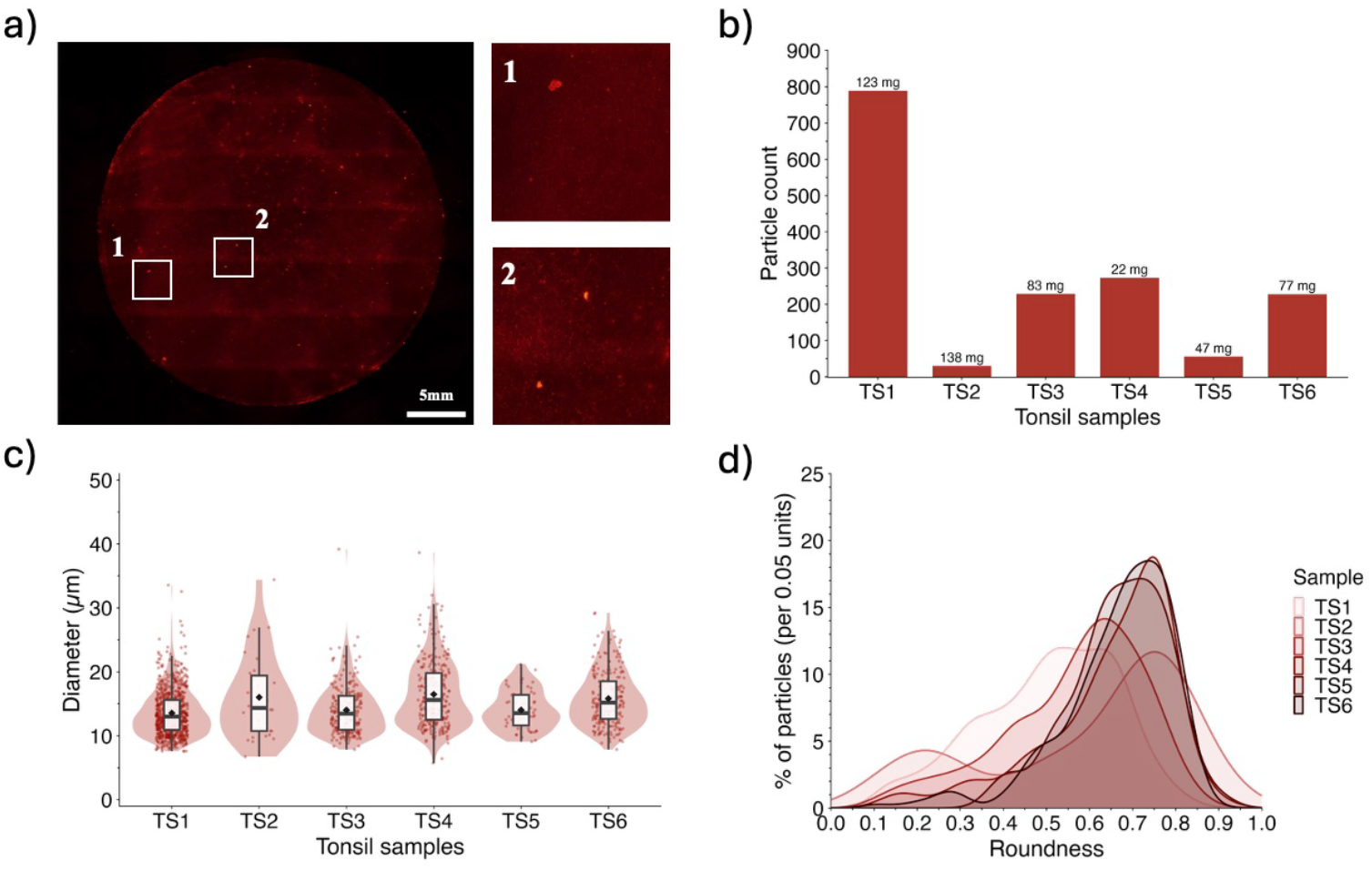
Nile Red fluorescence microscopy of tonsillar NMP particles. a) Representative image for microplastics filtered on a AlOx filter (4X magnification; Ex: 560 nm Em: 630 nm) b) Total particle count across all tonsil samples. c) Size distribution of all quantified microplastics d) Smoothed kernel-density estimates of particle roundness for samples TS1–TS6, rescaled to percentage per 0.05 roundness units and normalised per sample as an indicator of microplastics morphology across all filtered tonsil samples Roundness: 0 = elongated, 1 = circular.

Preliminary inspection of PCA loading patterns suggested that select polymers, including styrene-butadiene rubber (SBR), may contribute disproportionately to cluster separation, although these findings remain exploratory and will require confirmation in larger cohorts and formal loading analyses. To aid visualization of multivariate structure, 95% confidence ellipses were overlaid for each cluster. These findings support the feasibility of using NMP compositional profiling to identify distinct polymer signatures within biological tissues and provide a framework for future studies examining environmental, biological, and disease-associated determinants of polymer accumulation.

### Fluorescence microscopy confirms microplastic particles in tonsil tissue

To corroborate the Py-GC/MS findings, microplastic particles were isolated from a subset of tonsil specimens (n = 6) using a combined chemical and enzymatic digestion protocol. The digestion protocol effectively degraded all organic tissue while preserving MPs. Analyses identified fluorescently labeled particles in all six tonsil samples. The tissue-free negative control yielded 39 particles, representing a low background relative to sample counts and confirming that procedural contamination was minimal. The median microplastic count was 229, with particle numbers ranging from 30 to 789 across all digested tonsil samples. Inter-sample variability is consistent with the pilot nature of this study but could also underscore heterogeneity in personal microplastic exposure. No correlation was observed between tonsil mass and particle quantity. Particle morphology was characterized by roundness and particle diameter. The particles had a mean diameter of 15.0 µm ± 1.2 across samples. The smallest particle diameter identified was 5.7 µm, and the largest was 39.2 µm. The particle roundness ranged from 0.11 to 0.93, with an average of 0.61 ± 0.07, indicating moderate elongation and considerable inter- and intra-sample shape variability. Notably, this technique has limited sensitivity for particles below ∼5-10 μm.

### O-PTIR spectroscopy confirms intratissue NMP localization

To distinguish true tissue-embedded NMPs from potential processing contaminants, O-PTIR spectroscopy was performed on cryosectioned (10 μm) tonsil tissue mounted on CaF^2^ slides. O-PTIR enables label-free, non-destructive chemical identification at sub-micrometer spatial resolution, overcoming the diffraction limits of conventional FTIR.^18–20^ Multiple foci of polytetrafluoroethylene (PTFE) were identified within the tissue parenchyma, confirmed by the characteristic asymmetric CF^2^ stretching peak at 1,215 cm^−1^ (Fig. 4a). Spatial mapping of this spectral signature demonstrated discrete PTFE deposits embedded within the tissue architecture (Fig. 4b), confirming that these particles were not surface contaminants introduced during processing. This represents, to our knowledge, the first application of O-PTIR for NMP detection in human lymphoid tissue.

**Figure 4.**
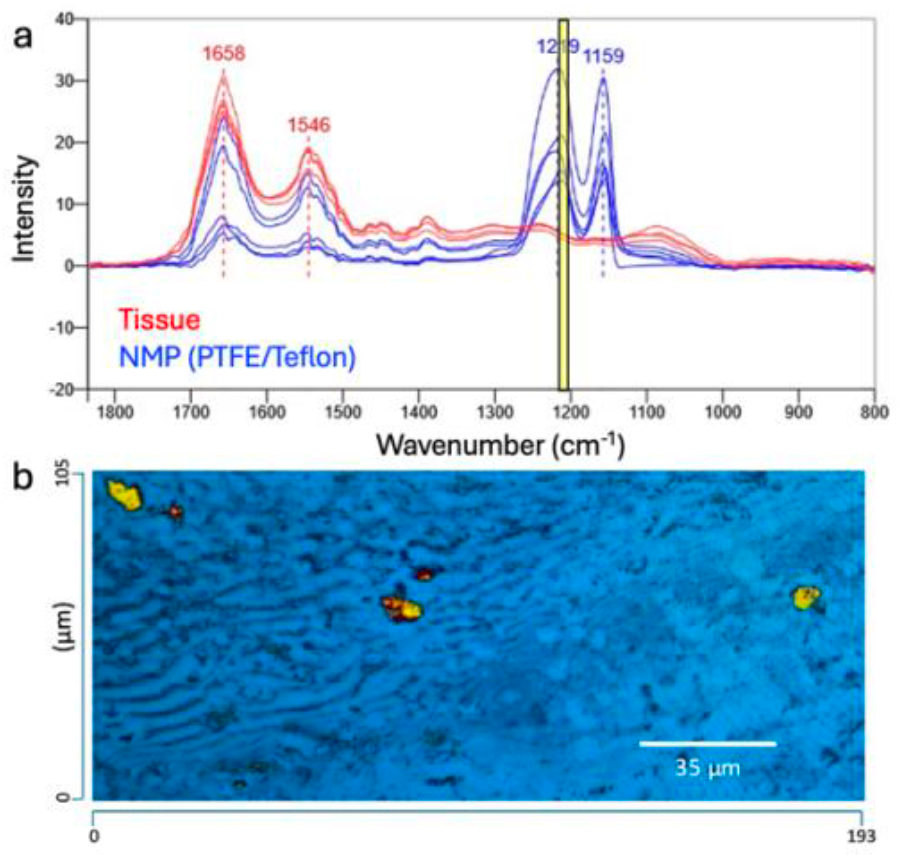
(a) OPTIR spectrum of areas with tissue (red) and PTFE NMPs (blue). Unique peaks at 1215 cm-1 (yellow) indicate asymmetrical stretching of CF2 which are unique to the NMP. (b) Mapping the spatial distribution of the 1215cm-1 peak (PTFE) in a tonsillar tissue section.

### Pilot dose–response studies reveal temporal and size-dependent immune responses

To investigate the immunomodulatory potential of NMPs, human lymphoid aggregate culture (HLAC) organoids derived from pediatric tonsil tissue^21^ were exposed to three microplastic conditions: PE 10 μm, PE 0.1 μm, and PS 0.35 μm, each at four concentrations (0, 1, 20, and 100 μg/mL) in triplicate. Supernatants were collected at day 3 and day 14 and analyzed using a Luminex 48-plex bead-based immunoassay (EMD Millipore) performed by the Human Immune Monitoring Center (HIMC) at Stanford University.

At day 3 post-exposure multiplex immunoassay data showed a significant upregulation of IL-6 (p = 0.011) and MIP-1β/CCL4 (p = 0.011), suggesting early innate immune activation with enhanced immune cell recruitment and potential skewing toward Th17 responses. By day 14, the cytokine landscape became more complex: low NMP concentrations were associated with decreased MDC/CCL22 (p 0.0001), suggesting compromised regulatory T-cell recruitment, while high concentrations (100 μg/mL) produced coordinated elevation of FLT-3L (p = 0.047), G-CSF (p = 0.047), and IL-4 (p = 0.047), indicating both tissue damage responses and potential type 2 immune reprogramming. Size-dependent effects were evident, with decreased IL-3 in organoids exposed to larger (10 μm) versus smaller (0.1 μm) particles (p = 0.015), suggesting differential cellular uptake or tissue architecture disruption as a function of particle size.

### Time-dependent NMP penetration into lymphoid organoids

To assess whether NMPs penetrate into the three-dimensional organoid architecture, HLAC organoids on transwell inserts were exposed to 100 nm fluorescent PE beads (100 μg/mL) and sectioned at day 3 and day 14. Z-binned fluorescence analysis of 10 μm cryosections demonstrated surface-enriched bead localization at day 3 (top > mid > bottom signal intensity). Quantitative line-based depth measurements (n = 12 per timepoint, 3 fields × 4 lines) revealed mean penetration depths of 57.5 ± 22.2 μm at day 3 versus 85.6 ± 26.1 μm at day 14 (Fig. 5a). After normalizing for tissue thickness, beads extended through 70% (mean = 0.70, SD = 0.14) of the organoid cross-section at day 3 versus 95% (mean = 0.95, SD = 0.02) at day 14 (Fig. 5b), suggesting time-dependent penetration independent of section thickness variation. Immunofluorescence co-staining with CD20 (B cells) and CD68 (macrophages/monocytes) demonstrated bead co-localization with both lymphocyte populations, with more extensive distribution among immune cell aggregates at day 14 compared to day 3.

**Figure 5:**
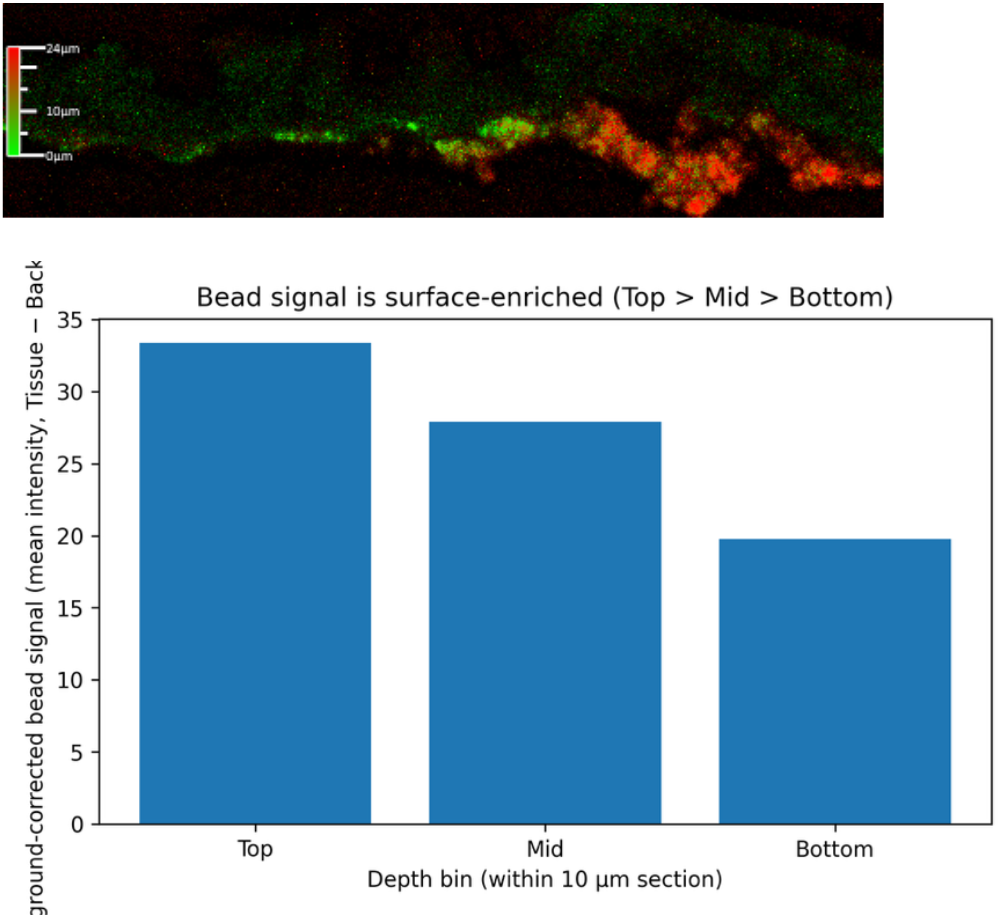
NMP penetration into tonsil organoids. A) Confocal microscopy of organoid cryosections showing 100 nm fluorescent PE bead distribution over time. B) Z-binned depth profiling quantifying bead signal in top, middle, and bottom depth bins. C) Line-based penetration measurements (n=12 per timepoint) from apical surface to deepest bead-positive signal, normalized to total tissue depth.

**Figure 6:**
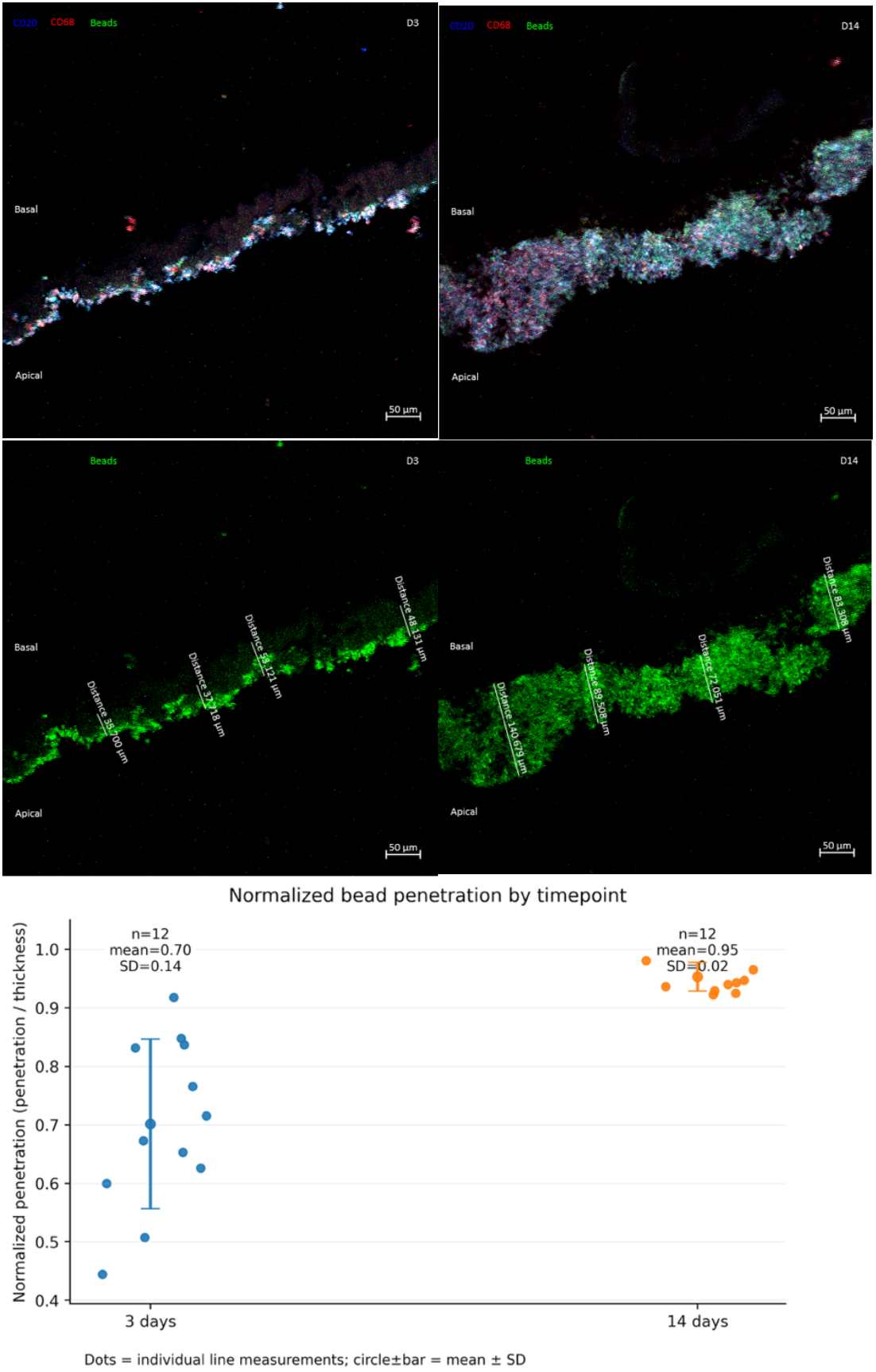
Cellular uptake and time-dependent distribution of NMPs within tonsil organoids. Immunofluorescence microscopy of tonsil organoid cryosections following exposure to fluorescent 100 nm PE beads. Top panel: CD20+ B cells (cyan) showing bead co-localization (magenta/red fluorescent beads) at day 3. Middle panel: CD68+ macrophages/monocytes (green) demonstrating bead uptake and distribution at day 14. Bottom panel: Quantification of normalized bead penetration depth by timepoint, showing individual measurements (dots; n=12 per timepoint from 3 fields × 4 lines) and mean ± SD. Beads penetrated through 70% (mean = 0.70, SD = 0.14) of organoid depth at day 3 versus 95% (mean = 0.95, SD = 0.02) at day 14, indicating time-dependent penetration into lymphoid tissue architecture. Bead co-localization was observed with both B cell and macrophage/monocyte populations, with more extensive distribution among immune cell aggregates at day 14 compared to day 3. Scale bars indicate magnification.

### Patient-derived multi-polymer challenge elicits coordinated immune module responses

Building on the tissue burden data, a cryo-milled multi-polymer mixture was formulated to reflect the pediatric tonsillar NMP profile (Figure 1): approximately 40% PE, 10% PET, 11% SBR, 10% N6, 10% N66, 6% ABS, 6% PVC, and 6% PP by mass (mean particle diameter 2.56 ± 1.45 μm), Figure 7. Some lower abundance polymers were not included in the HCM such as polystyrene^22^, poly(methyl methacrylate), and polycarbonate (PC). Source polymers were obtained from the University of Hawaii Polymer Kit and cryo-milled using a Retsch cryomill under liquid nitrogen cooling without direct particle contact at Oregon State University (SH).

**Figure 7:**
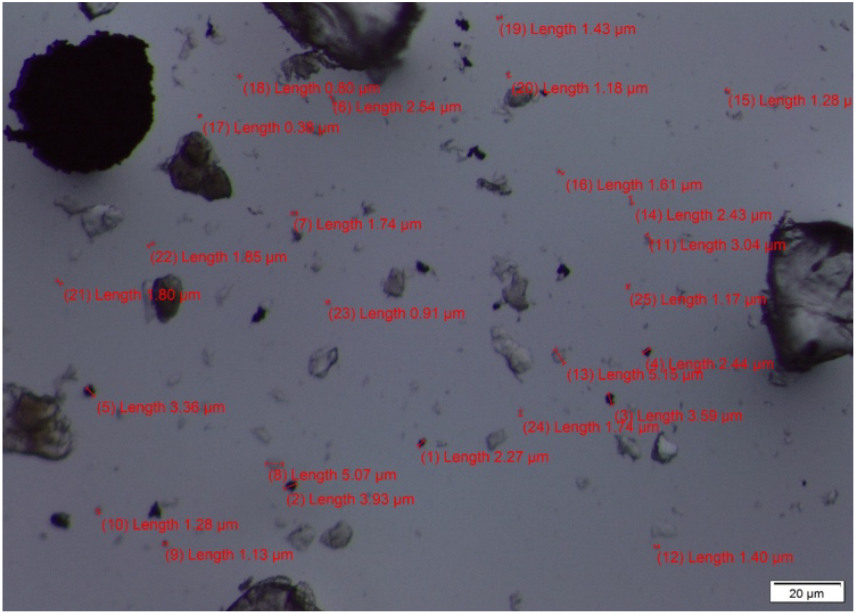
Human-composition multi-polymer mixture (HCM) formulated from pediatric tonsil burden data. Brightfield microscopy of cryo-milled multi-polymer mixture designed to reflect the NMP profile detected in pediatric tonsil tissue (Figure 1). Representative particle size measurements are indicated in red annotations, with mean particle diameter of 2.56 ± 1.45 μm. The mixture exhibits particle size heterogeneity and irregular morphology characteristic of environmentally weathered microplastics. Scale bar = 20 μm.

We employed a vascularized HLAC tonsil organoid (ETV2 iPSC + HLAC) model developed in our lab so that we would be able to extend the viability of the organoid model to support that the change in data was secondary to the response to the exposure and not an artifact of the organoid model degrading. Vascularized HLAC tonsil organoids (n = 5 per condition) were exposed to control medium, 100 nm PE, or the human-composition mixture (HCM) with longitudinal supernatant collection at days 3, 7, 11, and 14. Non-vascularized organoids (n = 12 per condition) were exposed to control or HCM with endpoint sampling at day 14. Supernatants (84 samples total) were analyzed using a Luminex 115-plex immunoassay.

Volcano plot analysis demonstrated robust early responses: at day 3, PE exposure produced 60 significant cytokine changes and HCM produced 85 significant changes (q 0.05) versus control. By day 14, the PE volcano was essentially empty (1 significant hit) while HCM retained 23 sIgnificant hits, indicating progressive convergence toward control levels (Fig. 8).

**Figure 8:**
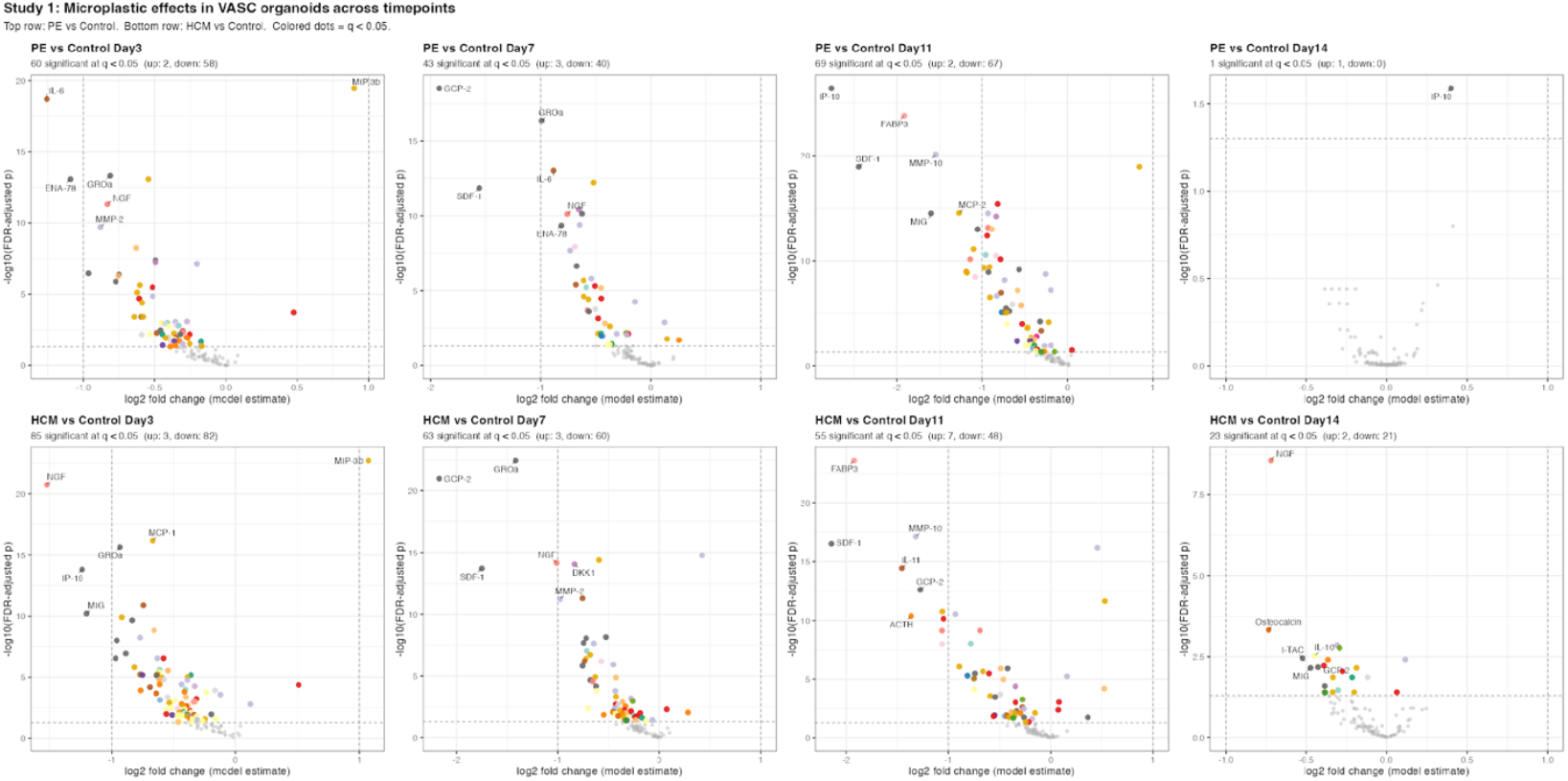
Temporal dynamics of NMP-induced cytokine responses in vascularized tonsil organoids. Volcano plot time series showing cytokine secretion profiles from vascularized HLAC tonsil organoids (ETV2 iPSC + HLAC; n=5 per condition) exposed to 100 nm PE (top row) or human-composition mixture (HCM, bottom row) versus control at days 3, 7, 11, and 14. Supernatants were analyzed using Luminex 115-plex immunoassay. X-axis shows log2 fold change (model estimate) relative to control; Y-axis shows -log10(FDR-adjusted p-value). Horizontal dashed line indicates significance threshold (q < 0.05). Colored dots represent significant cytokines (orange/yellow/red scale), with select cytokines labeled. At day 3, PE exposure produced 60 significant cytokine changes and HCM produced 85 significant changes versus control, demonstrating robust early immunomodulatory responses. By day 14, the PE condition showed only 1 significant hit while HCM retained 23 significant hits, indicating progressive convergence toward control cytokine levels over time. This temporal pattern suggests dynamic immune adaptation in response to NMP exposure in organized lymphoid tissue.

**Figure 9:**
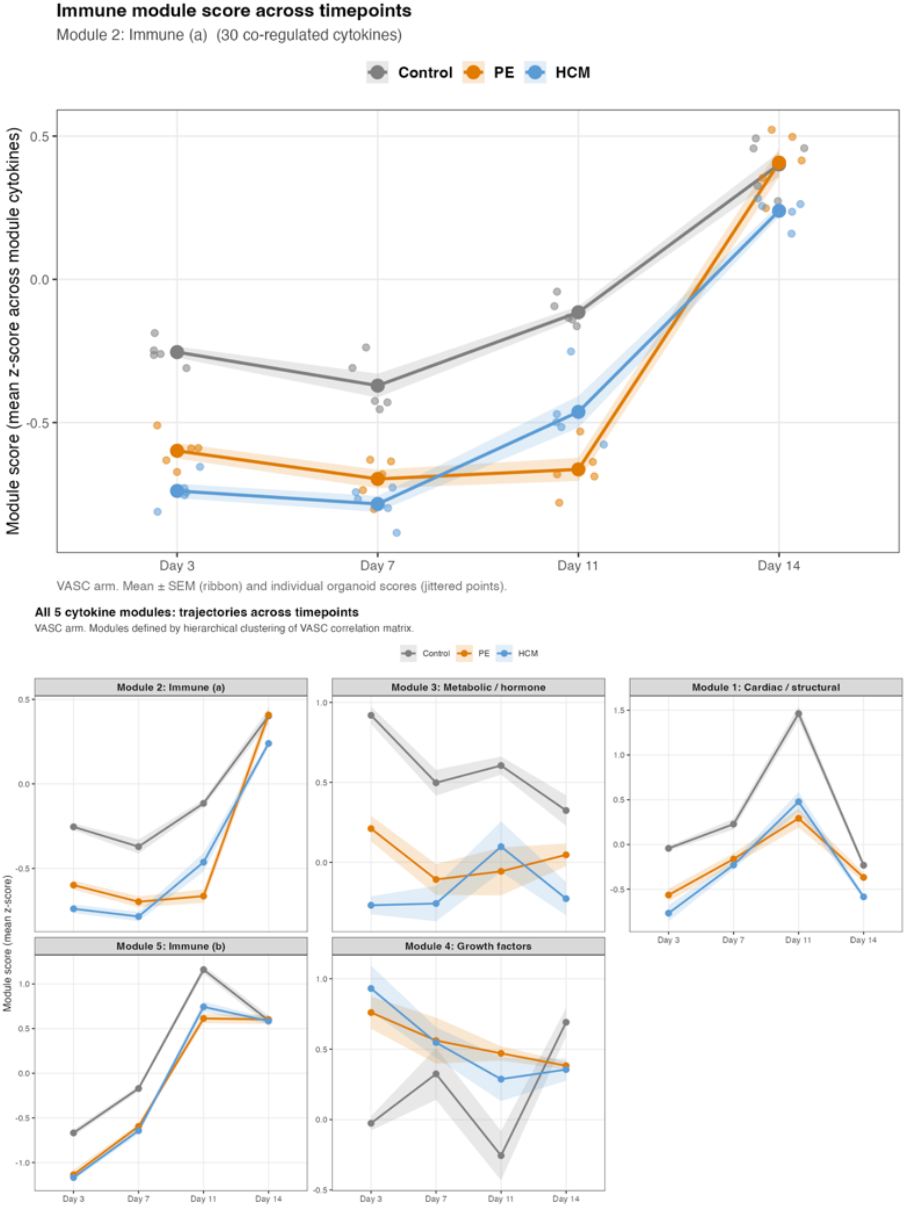
Cytokine module trajectory analysis reveals coordinated temporal responses to NMP exposure. Top panel: Representative trajectory plot for Module 2: Immune (a), containing 30 co-regulated cytokines. Mean z-score (mean ± SEM ribbon) and individual organoid scores (jittered points) are shown for Control (gray), 100 nm PE (orange), and HCM (blue) conditions across days 3, 7, 11, and 14 in vascularized organoids (n=5 per condition). Both PE and HCM conditions show maximal deviation from control at day 3, followed by progressive convergence toward control levels by day 14. Bottom panel: Trajectory plots for all five cytokine modules identified by hierarchical clustering of the 115-analyte correlation matrix from vascularized organoids. Modules include: Module 2: Immune (a), Module 3: Metabolic/hormone, Module 1: Cardiac/structural, Module 5: Immune (b), and Module 4: Growth factors. Lines show mean module z-scores with shaded ribbons representing SEM. All five modules demonstrate a consistent pattern of maximal deviation at day 3 with progressive convergence by day 14 for both PE and HCM exposures. Importantly, responses within each module are bidirectional (not uniformly upregulated or downregulated), arguing against non-specific artifacts and supporting genuine cytokine profile remodeling in response to NMP exposure. This coordinated temporal pattern suggests organized immune adaptation across multiple functional pathways in lymphoid tissue.

**Figure 10:**
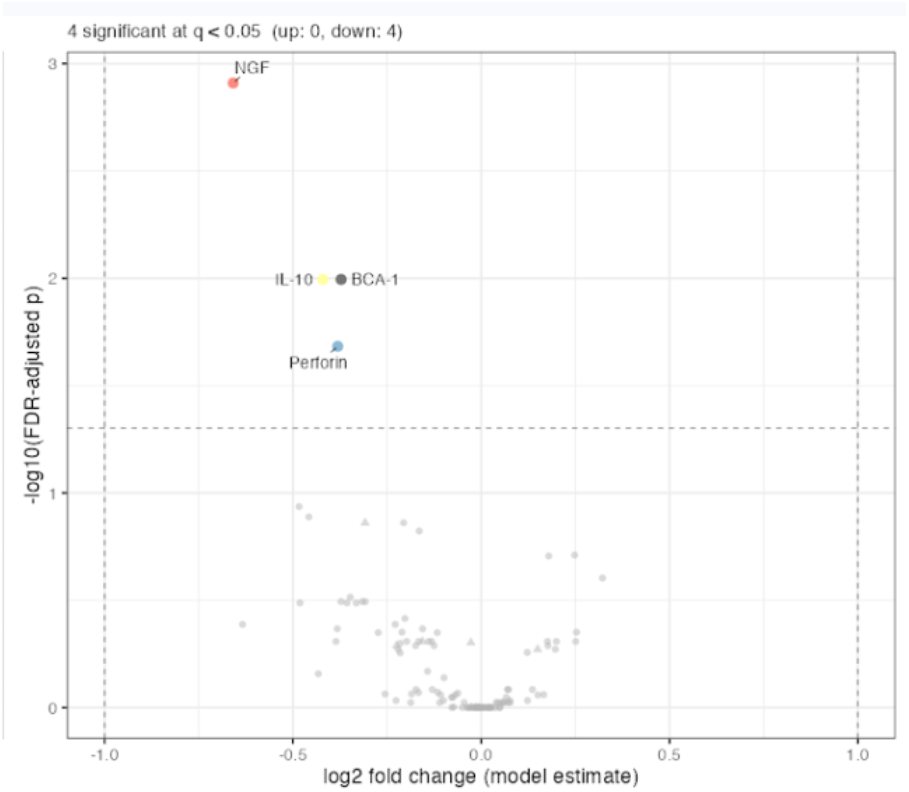
Vascularization has minimal impact on NMP-induced cytokine responses. Volcano plot comparing cytokine secretion profiles between vascularized and non-vascularized tonsil organoids at day 14 following NMP exposure (115-plex Luminex analysis). X-axis shows log2 fold change (model estimate) between vascularized and non-vascularized conditions; Y-axis shows -log10(FDR-adjusted p-value). Horizontal dashed line indicates significance threshold (q < 0.05). Only 4 of 115 cytokines showed significant differential expression: NGF (upregulated), and IL-10, BCA-1, and Perforin (downregulated) in vascularized versus non-vascularized organoids.

Hierarchical clustering of the 115-analyte dataset identified five co-regulated cytokine modules: immune (a), immune (b), metabolic/hormone, cardiac/structural, and growth factors. Module trajectory analysis revealed a consistent pattern across all five modules: both PE and HCM conditions deviated maximally from control at day 3, with progressive convergence by day 14 (Fig. 8). Importantly, these changes were bidirectional within each module—not uniformly upregulated or downregulated—arguing against a non-specific artifact and supporting cytokine profile remodeling.

Comparison of vascularized versus non-vascularized organoids at day 14 revealed that only 4 of 115 cytokines showed differential effects attributable to the presence of endothelial cells, suggesting that the immunomodulatory response to NMPs may be driven by immune and stromal cell populations rather than the vascular compartment.

## Methods

### Study design and tissue collection

Pediatric tonsil specimens were collected from children undergoing tonsillectomy for standard clinical indications (obstructive sleep apnea and/or recurrent tonsillitis) under an institutional review board–approved protocol. Specimens were collected using metal instruments, immediately wrapped in sterile aluminum foil in the operating room, weighed, and stored in a steel box in -80degree freezer to minimize ambient contamination. Tissue was collected at a single institution under a single low-contamination protocol. Care was taken in subsequent analyses and tissue manipulation to avoid processing contamination including working under laminar flow hoods, rinsing with filtered solvents, and minimizing plastic labware and clothing to the extent possible.

### Py-GC/MS polymer analysis

Py-GC/MS was performed at the University of New Mexico (MC, RT). Tonsil tissue was prepared using three parallel methods: (1) ethyl acetate/methanol extraction, (2) KOH digestion with ultracentrifugation, and (3) KOH digestion with cyclohexane filtration. Polymer identification was performed by Py-GC/MS with a >65% probability threshold. Unsupervised analysis included PCA and K-means clustering (k = 2) of polymer concentration profiles.

### Nile Red fluorescence microscopy

Fluorescence microscopy was performed at Indiana University (NA, AB). Tissue (n = 6) underwent sequential chemical digestion (10% KOH, 50°C, 48 h, 42 kHz sonication) and enzymatic digestion (DNase 100 μg/mL, collagenase 1 mg/mL, 37°C, 2 h; proteinase K 250 μg/mL, 50°C, 24 h). Digests were adjusted to 5 M urea, stained with Nile Red (30 μg/mL, 70°C, 8 min), filtered through 0.2 μm aluminium oxide filters using an all-glass vacuum apparatus to minimize exogenous plastic contamination, and imaged on a Nikon Eclipse Ni-L microscope (TxRed ex-560 nm, Cy3 ex-540 nm). From these images, particle count, diameter, and area were quantified using R statistical software. Particle shape (roundness) was derived from diameter measurements. A tissue-free negative control was processed in parallel to assess background contamination.

### O-PTIR spectroscopy

Tonsil tissue was cryosectioned at 10 μm and mounted on CaF^2^ slides. O-PTIR spectra were acquired to identify polymer-specific absorption peaks, with spatial mapping of the 1,215 cm^−1^ PTFE signature.^18,20^

### HLAC tonsil organoid culture

Human lymphoid aggregate culture organoids were generated from freshly excised pediatric tonsil tissue as previously described. For vascularized models, endothelial cells were incorporated to extend organoid viability over the 14-day experimental window. Non-vascularized organoids (∼6 million cells; ∼0.9 mg dry mass) were cultured on transwell inserts.

### NMP exposure conditions

Pilot studies used commercial PE (10 μm and 0.1 μm) and PS (0.35 μm) particles at 0, 1, 20, and 100 μg/mL. Expanded studies used cryo-milled HCM (40% PE, 10% PET, 11% SBR, 10% N6, 10% N66, 6% ABS, 6% PVC, 6% PP; mean diameter 2.56 ± 1.45 μm) and 100 nm PE, with doses scaled to organoid dry mass (a tonsil organoid is ∼6 million cells and measures ∼0.9 mg dry mass) based on pediatric tonsil tissue burden data. For the HCM, plastics were received from the Hawaii Polymer Kit produced by the University of Hawaii. Samples were cryomilled using a Retch cryomill where plastics are ball milled while the chamber is bathed in liquid nitrogen (no direct liquid nitrogen contact with particles). The resulting cryomilled plastics were assessed visually for particle sizes using a dissecting scope equipped with a camera, such that four images were taken, and 25 particles were randomly chosen and measured per image for a total of 100 particles assessed for each polymer. Following individual polymer size distribution assessment, each polymer was mixed in the percentages provided and sent to Kara Meister at Stanford University.

### Luminex cytokine profiling

Culture supernatants were analyzed by Luminex bead-based sandwich immunoassay (48-plex, EMD Millipore, for pilot studies; 115-plex for expanded studies) performed by the HIMC at Stanford University. Median fluorescence intensity (MFI) data were log^2^(MFI + 1) transformed. For the pilot 48-plex data, quantile regression was used to model the median, with MFI data centered, scaled, and regressed on block, microplastic type, concentration, and log(CHEX4 MFI) as a covariate for non-specific binding; p-values were FDR-adjusted at 5% across proteins. For the expanded 115-plex longitudinal data, linear mixed-effects models were used for the vascularized arm (with organoid as random effect) and linear models for the day 14 cross-arm comparison. Cytokine modules were defined by hierarchical clustering of the vascularized-arm correlation matrix. Analyses were performed in R v.4.5.3 using tidyverse, lme4, lmerTest, emmeans, vegan, uwot, ComplexHeatmap, ggplot2, ggrepel, and patchwork.

### Organoid penetration analysis

Organoids exposed to 100 nm fluorescent PE beads were cryosectioned and imaged by confocal microscopy. Z-binned depth profiling quantified bead signal in top, middle, and bottom depth bins. Line-based penetration measurements (n = 12 per timepoint) were performed from the apical surface to the deepest bead-positive signal, normalized to total tissue depth.

### Transmission electron microscopy

Organoids exposed to 100 nm PE (100 μg/mL, 14 days) were fixed, embedded, and sectioned for TEM. PE presence was confirmed by fluorescence prior to processing.

### Statistical analysis

All statistical analyses were performed in R v.4.5.1 using packages including openxlsx, Matrix, and quantreg. Multiple comparison corrections used FDR at 5%. PCA and K-means clustering were performed on centred and scaled polymer concentration data.

## Discussion

This study provides three principal advances in understanding the relationship between environmental plastic contamination and pediatric immune health. First, it establishes that NMPs are ubiquitous in pediatric tonsil tissue, with a heterogeneous polymer composition reflecting the dual inhalation and ingestion exposure routes of the aerodigestive tract. Second, it demonstrates that NMPs are embedded within the tissue parenchyma and not merely surface contaminants by using O-PTIR spectroscopy, a novel technique recognized for the capacity to detect and characterize plastics at sub-micrometer resolution in intact biological specimens.^19,20^ Third, it shows that environmentally relevant NMP mixtures, formulated to match the patient-derived polymer profile, elicit significant but transient immunomodulatory responses in human tonsil organoids following an acute exposure.

The detection of NMPs in all tested pediatric tonsil specimens is consistent with the emerging picture of widespread human tissue contamination. Zhu et al. reported microplastic concentrations of 6.03 ± 7.37 particles/g in adult tonsil tissue using laser direct infrared spectroscopy, and Nihart et al. demonstrated increasing NMP concentrations in human brain tissue over time.^6,23^ The polymer profile observed here was dominated by PE, PET, and ABS. This partially overlaps with but is distinct from profiles reported in brain (PE-dominant) and blood (PE, PP, PS predominant), suggesting tissue-specific accumulation patterns that may reflect local exposure routes, particle size-dependent filtration, or differential polymer–tissue interactions.^24^ The field is also new and different investigations have used different methods.^25^ The distinct polymer profile found in this study could also be due to differences in methods, preparation (e.g. digestion) techniques of previous studies or the probability cutoffs used rather than truly different accumulation patterns by tissue type.

The tonsil represents a particularly informative model for studying NMP–immune interactions. As a secondary lymphoid organ with organized germinal centers, crypt epithelium facilitating antigen sampling, and dense populations of B cells, T cells, macrophages, and dendritic cells, the tonsil recapitulates the full spectrum of mucosal immune responses. The observation that NMPs are embedded within the tissue raises the possibility that chronic particle exposure contributes to the inflammatory remodeling, crypt distortion, and lymphoid hyperplasia that characterize common pediatric tonsillar disease phenotypes.

The cytokine profiling data propose several possible immunologic responses, all which warrant further investigation. The early (day 3) upregulation of IL-6 and MIP-1β/CCL4 is consistent with innate immune activation and monocyte/macrophage recruitment, aligning with prior reports of IL-6 elevation in response to microplastic exposure in liver organoids and primary human immune cells.^13,26^ The subsequent decrease in MDC/CCL22 at low concentrations suggests potential disruption of regulatory T-cell recruitment, while the coordinated elevation of FLT-3L, G-CSF, and IL-4 at high concentrations could favor stress responses and type 2 immune skewing. These findings are consistent with the “dualism” of pro- and anti-inflammatory responses to microplastics described by Wolff et al., who observed simultaneous IL-17A elevation and IL-1β suppression in primary human immune cells exposed to PS and PMMA particles.^13^

The convergence of treated and control cytokine profiles by day 14 in the vascularized organoid model is also an observation which warrants further investigation. The temporal pattern of maximal deviation at day 3 with progressive normalization was consistent across five functional cytokine modules and both PE and HCM conditions. This may reflect immune adaptation, particle clearance or sequestration, or the inherent resilience of organized lymphoid tissue. However, it does not exclude the possibility that repeated or chronic exposure as would occur *in vivo* could sustain, amplify, or otherwise alter these responses from physiological adaptation toward pathological remodeling.

The time-dependent penetration of nanoscale particles into organoid aggregates (70% depth at day 3 to 95% at day 14) and their intracellular localization within lymphocyte lysosomes on TEM provide mechanistic evidence for active NMP uptake by immune cells. This is consistent with the phagocytic uptake of NMPs by macrophages and dendritic cells described in vitro and the observation of NMP deposition in cerebrovascular immune cells in human brain tissue.^6,13^ The mechanisms underlying these findings will require further study.

A distinguishing feature of this study is the translational paradigm of using clinical tissue burden data to directly inform preclinical exposure conditions. The cryo-milled multi-polymer mixture was formulated to match the polymer composition detected in patient tonsils, and exposure concentrations were scaled to organoid dry mass. This approach addresses a major limitation of existing NMP immunotoxicity studies, which typically use single commercial polymers (most commonly PS) at concentrations that may not reflect real-world exposures.^4,12^ The finding that the multi-polymer HCM produced a more sustained immune response than single-polymer PE (23 versus 1 significant cytokine changes at day 14) underscores the importance of studying NMPs as complex environmental mixtures.

## Limitations

Several limitations warrant consideration. The sample size (n = 30 for Py-GC/MS, n = 6 for fluorescence microscopy) limits statistical power for clinical correlations. There is likely an underestimate of nanoscale polymers in the data generated by fluorescence microscopy because of the technical limitations of that approach. The organoid penetration data are from single-donor pilot experiments requiring replication. The Luminex assay measures secreted cytokines in culture supernatant and does not provide single-cell resolution of which immune cell populations are responsible for specific cytokine changes. The 14-day observation window, while extended relative to most in vitro studies, cannot capture the chronic exposure dynamics relevant to *in vivo* tonsillar disease. Contamination controls, while implemented (blank samples, metal instruments, aluminum oxide filters), cannot entirely exclude background NMP exposure during processing. Finally, the clinical significance of the observed cytokine changes and whether they translate to meaningful immune dysfunction in children remains to be established. Future studies teasing out the mechanisms driving the observed preliminary findings reported here will be necessary to further understand and characterize the clinical burden of these results.

## Conclusion

This study demonstrates that NMPs are universally present in pediatric tonsil tissue, are embedded within the lymphoid parenchyma, and elicit significant immunomodulatory responses when applied to human tonsil organoids at environmentally relevant compositions and concentrations. By establishing a translational pipeline from clinical tissue characterization to patient-informed preclinical modeling, this work provides a framework for investigating NMP–immune interactions in physiologically relevant human systems. The findings have implications for understanding the rising prevalence of immune-related disorders in children and for informing evidence-based public health strategies to mitigate pediatric NMP exposure.

## Data availability

All data supporting the findings of this study are available from the corresponding author upon reasonable request. Source data for figures are provided with this paper.

## Acknowledgements

The authors acknowledge the Human Immune Monitoring Center (HIMC) at Stanford University for performing the Luminex immunoassays.

Funding from the National Science Foundation Growing Convergence Research [Grant 1935028] to SH.

Support from The Stanford Woods Institute for the Environment and the Center for Human and Planetary Health Early Career Award to KM.

This project/publication was additionally supported by the Stanford Maternal and Child Health Research Institute, the Department of Otolaryngology—Head & Neck Surgery at Stanford University School of Medicine, and philanthropic support to KM.

